# MGTdb: A web service and database for studying the global and local genomic epidemiology of bacterial pathogens

**DOI:** 10.1101/2022.06.14.496187

**Authors:** Sandeep Kaur, Michael Payne, Lijuan Luo, Sophie Octavia, Mark M. Tanaka, Vitali Sintchenko, Ruiting Lan

## Abstract

Multilevel genome typing (MGT) enables the genomic characterization of bacterial isolates and the relationships among them. The MGT system describes an isolate using multiple multilocus sequence typing (MLST) schemes, referred to as levels. Thus, for a new isolate, sequence types (STs) assigned at multiple precisely defined levels can be used to type isolates at multiple resolutions. The MGT designation for isolates is stable, and assignment is faster than existing approaches. MGT’s utility has been demonstrated in multiple species.

This paper presents a publicly accessible web service called MGTdb, which enables the assignment of MGT sequence types to isolates, along with their storage, retrieval and analysis. The MGTdb web service enables upload of genome data as sequence reads or alleles, which are processed and assigned MGT identifiers. Additionally, any newly sequenced isolates deposited in NCBI Sequence Read Archive are also regularly retrieved (currently daily), processed, assigned MGT and made publicly available in MGTdb. Interactive visualisation tools are presented to assist analysis, along with capabilities to download publicly available isolates and assignments for use with external software.

MGTdb is currently available for *Salmonella enterica* serovars Typhimurium and Enteritidis, and *Vibrio cholerae*. We demonstrate the usability of MGTdb through three case studies to study the long-term national surveillance of *S*. Typhimurium, and the local epidemiology and outbreaks of *S*. Typhimurium, and the global epidemiology of *V. cholerae*. Thus, MGTdb enables epidemiological and microbiological investigations at multiple levels of resolution for all publicly available isolates of these pathogens.

**Database URL:** https://mgtdb.unsw.edu.au

## 1. Introduction

To prevent the spread of infectious diseases within populations, public health surveillance programs are implemented to enable early detection of outbreaks and epidemics, and monitoring and evaluation of disease prevention and control programmes (1). For diseases with epidemic or outbreak potential communicated by bacteria, such as salmonellosis or cholera, whole genome sequencing (WGS) of the causative pathogens provides high resolution epidemiological surveillance and tracing (2, 3). Hence, all modern surveillance systems use genomics to characterize (or type) isolates as well as to study the relationships between them (4). Additionally, adoption of WGS by public health laboratories both national and international (such as Public Health England (5) and PulseNet International (6)), continues to facilitate research and investigation into WGS-based methods that are feasible for epidemiological studies given the rapid accumulation of sequence data.

From whole genome sequences, the traditional computational method to investigate genetic relationships has been to use single nucleotide polymorphisms (SNP) (calculated via alignment to a reference sequence) to first build phylogenetic trees, followed by using that tree to infer the relationships among the isolates, such as determining if certain isolates share a common source (7). The use of this approach has been extensively employed for epidemiological tracing (8). For example, Ford *et al*. (9) linked seven *Salmonella* outbreaks to a common source, using maximum likelihood core genome SNP phylogeny. Phylogenetic analysis is considered the gold standard in establishing relatedness between isolates, as this approach can use information contained within the genome and provide detailed divergence information (10). However, this approach is limited in scalability, as the phylogeny often needs to be recomputed for each new isolate that becomes available, which is a computationally intensive and time-consuming task especially for a large number of genomes (10). This problem hinders continuous surveillance systems. Although computationally less-intensive approaches exist, such as alignment-free phylogenetics (11), such approaches are not commonly used in surveillance systems as their utility for bacterial source tracing and outbreak detection (short-term epidemiology) is not well demonstrated (12).

Most surveillance systems implement multilocus sequence typing (MLST) based methods (13) for characterising isolates. In this approach, when a new isolate becomes available for analysis, alleles are determined at a selected set of loci and assigned numeric identifiers. Then, the ordered combination of alleles, termed the allelic profile, is assigned a unique numeric identifier termed the sequence type (ST). Thus, an isolate is uniquely identified by its assigned ST, and two isolates with the same ST are considered identical. To infer relationships between isolates which are non-identical, a pairwise allele distance matrix is calculated, and single linkage clustering is performed at various levels of allele differences. The resulting clusters are termed clonal complexes (CC). Several MLST schemes have been developed enabling the analysis of isolates at varying levels of relatedness. A few commonly used schemes are the seven-gene MLST - comprising a set of seven housekeeping genes, ribosomal MLST (rMLST) - comprising the necessarily present and conserved set of ribosomal protein coding genes, core genome MLST (cgMLST) - comprising the core loci set of a species and whole genome MLST (wgMLST) - comprising all genes present in a species as well as intergenic regions. Such schemes are widely used to study isolates both globally and locally. For example, Schjørring *et al*. (14) used whole genome sequences to calculate cgMLST STs, and then a minimal spanning tree with a maximal allowed difference of five alleles for grouping together isolates, to trace the source of *Listeria* infections in Denmark - thus exemplifying the use of cgMLST to study local epidemology. In 2018, Alikhan *et al*. (15) analysed the population structure of *Salmonella* at varied levels of resolution using seven-gene MLST, rMLST, cgMLST and wgMLST for >110,000 *Salmonella* genomes, recommending cgMLST for epidemiological investigations. Enterobase (16), the web-service providing cgMLST for *Salmonella* and other species, implements hierCC (17), wherein cgMLST-CCs are calculated and presented at multiple levels of allele differences. Here, CCs calculated at larger levels of allele differences are used to study distantly related isolates, and CCs calculated at smaller levels of allele differences are used to study closely related isolates - thus providing a strategy for studying relationships between isolates at varying levels of resolution using a single cgMLST scheme.

Allele based typing followed by single linkage-clustering, is preferred due to its portability and standardized-nomenclature compared to SNP-based phylogenetic approaches (18). The concept of allele typing has also been implemented directly at the SNP-level - as exemplified by the SNP address implemented in SnapperDB (19). Uelze *et al*. (20) provide a summary of WGS based typing methods and computational systems which implement these approaches. **Table 1**, summarizes the commonly used web-systems for epidemiological analysis and surveillance, and the methods they implement.

**Table 1.** Commonly used web-based systems incorporating genomics based methods for surveillance of pathogenic bacteria.

| **Tool** | **Availability** | **Method** | **Application species** |
| --- | --- | --- | --- |
| Enterobase (16) | Publicly available online<br>(https://enterobase.warwick.ac.uk) | MLST, cgMLST, hierCC | 8 organisms |
| SnapperDB (19) | Code available online<br>(https://github.com/phe-bioinformatics/snapperdb) | SNPs calculated w.r.t. a reference, linkage clustering at 7 thresholds. | Used by Public Health England for *Salmonella*. |
| SeqSphere+ | Commercial<br>(https://www.ridom.de/seqsphere/) | MLST, cgMLST, linkage clustering and more | 54 organisms |
| BacWGSTdb (31) | Publicly available online<br>(http://bacdb.cn/BacWGSTdb/) | MLST, cgMLST, SNP phylogenetic trees and minimal spanning tree. | 20 organisms |
| PathogenWatch (32) | Publicly available online<br>(https://pathogen.watch) | MLST, cgMLST, single linkage clustering and coreSNP phylogenetic trees. | 13 organisms |
| PubMLST (33) | Publicly available online<br>(https://pubmlst.org) | MLST based schemes, linkage clustering | 130 organisms |

All approaches which rely on linkage-clustering to determine isolate relationships have two major disadvantages 1) the clusters need to be recomputed for each new isolate, 2) a new isolate may be equidistant to two (or more) clusters. In the latter case, the clusters may merge and hence the nomenclature used to refer to related isolates may not be stable in all cases. An approach to provide naming stability is to artificially prevent the two clusters from merging and to arbitrarily assigned the new isolate to a cluster (e.g. in HierCC (17), the new isolate is assigned to the cluster with the smaller label). In such approaches, different orders of strain addition can affect clustering results from the same set of isolates and true genetic relationships may not necessarily be reflected in the inferred clusters.

Building on the well accepted MLST-based methods, our group previously developed multilevel genome typing (MGT) (21), a typing system consisting of multiple ordered MLST schemes (referred to as MGT levels) within the single approach. The MGT levels have been set up such that the number of loci, and therefore, the likelihood of observing a mutation (a new allele) in each level, increases from MGT1 onwards - thus lower levels offer lower resolution and larger levels offer higher resolution. Using the MGT approach, each isolate is assigned a sequence type (ST) at each MGT level. The isolate’s MGT identity is then defined as the series of STs at each level, in ascending MGT-level order (referred to as the genome type (GT)). The STs and GTs are stable and will not change with the addition of new isolates. MGT levels differ from levels of clusters defined using hierarchical clustering as they are stable. Furthermore, calculating STs is faster than calculating CCs, as the computation of an allele distance matrix is not required. Along with the stable STs and GTs, MGT also assigns CCs and outbreak detection clusters (ODC). The CCs defined here group STs by single linkage clustering (22) with a threshold of at most one allele difference, and ODCs group the largest MGT-level STs, allowing a defined number of allele differences. The maximal number of allele differences allowed by an ODC is identifiable from the number following the ODC - for example, ODC5 is defined using single linkage clustering with a cut-off of at most five allele differences. For all species that have been described using MGT so far (21, 23, 24), four levels of ODC have been defined (ODC1, ODC2, ODC5 and ODC10). The CCs and STs are only calculated for further, or alternative, relationship analysis between isolates and are not recommended for relationship characterisation. MGT has been developed and successfully demonstrated for studying the global and local epidemiology of *Salmonella enterica* serovars Typhimurium (21) and Enteritidis (23) and *Vibrio cholerae* (24), comprising nine, nine and eight MGT levels, respectively.

In this paper, we present MGTdb (https://mgtdb.unsw.edu.au), a web-based service that enables a user to upload their isolates for assignment of MGT, along with storage, retrieval and analysis of these isolates at both global and local scales. The three pathogens previously successfully described using MGT (21, 23, 24) are available on MGTdb. **Section 2** describes the various features of the MGTdb web service. In **Section 3**, the usability of MGTdb is presented through three case studies and finally **Section 4** summarises and concludes this work.

## 2. The MGTdb web service

### 2.1 Software

The webservice is available at https://mgtdb.unsw.edu.au. Documentation is available at https://mgt-docs.readthedocs.io/en/latest/. A locally installable version of the software is available on github at https://github.com/LanLab/MGT.

### 2.2 Implementation

The MGTdb web application has been built in python (https://www.python.org; v3.7) using the django framework (https://www.djangoproject.com; v3.1), and is served via an Apache HTTP server (https://httpd.apache.org; v2.4) using mod_wsgi (https://modwsgi.readthedocs.io/en/master/; v4.6). The data associated with each target bacterial species is stored in a bacteria-specific PostgreSQL database (https://www.postgresql.org; v10.17) (for schema see **Supplementary Figure S1**). An additional PostgreSQL database stores user-login information. The client-side visualisations have been built using javascript’s D3 library (https://d3js.org; v6).

### 2.3 Workflow

A user can upload an isolate as Illumina paired-end reads, from which a genome is assembled and alleles extracted on the server. Alternatively, the user can upload allele sequences directly - we recommend using our previously published pipeline to extract the alleles locally (https://mgt-docs.readthedocs.io/en/latest/local_allele_calling.html#local-allele-calling) (21). As the file size of allele sequences is very small compared to raw reads, the latter method allows a user to upload multiple isolates at once. Any new alleles will be assigned a new unique identifier. An isolate will then be assigned a MGT by assigning STs (by assignment of identifiers to allelic profiles) and CCs (assignment of identifiers to single linkage clusters) for all MGT levels, and ODCs (assignment of identifiers to linkage clusters) for the largest MGT level (**Supplementary Figure S2**).

### 2.4 Database population

Apart from user-uploaded genome sequences of new isolates, any new whole-genome-sequenced isolates deposited in NCBI-SRA (25) for each hosted organism, are retrieved daily, passed through the MGT assignment pipeline, assigned MGT associated identifiers and are made available publicly on the MGTdb web-server (**Supplementary Figure S2**).

### 2.5 Search

Isolates in MGTdb can be searched using any MGT associated identifiers, as well as any metadata fields such as location and time. Currently, for simplicity, a user can choose to perform either union (OR) or intersection (AND; default selection) search, between all searched terms within a search.

### 2.6 Privacy

Due to privacy and security concerns, a user can upload their genome sequence as ‘private’, which disables the submitted isolate (and associated metadata) from being publicly searched or viewed, i.e. such an isolate can be viewed and searched only by the authorised user who uploaded it. A user can delete their uploaded-isolates (and associated metadata) at any time. While a user may hesitate to upload full genome sequences with associated accessory genome information such as antimicrobial resistance genes, the MGTdb web service alleviates these concerns by allowing only information required for MGT assignment to be uploaded, i.e. the allele sequences and minimum metadata (year and country). Furthermore, if a user does upload the whole genome sequence of an isolate, only the extracted alleles are retained, and the uploaded file is immediately deleted following the extraction of alleles.

### 2.7 Data visualisation and download

To enable effective exploration of data, a number of interactive and static visualisations are made available within MGTdb (summarized in **Figure 1**). These features are briefly described below; a detailed description of each of these features is presented in **Supplementary Note S1**.

**Figure 1.**
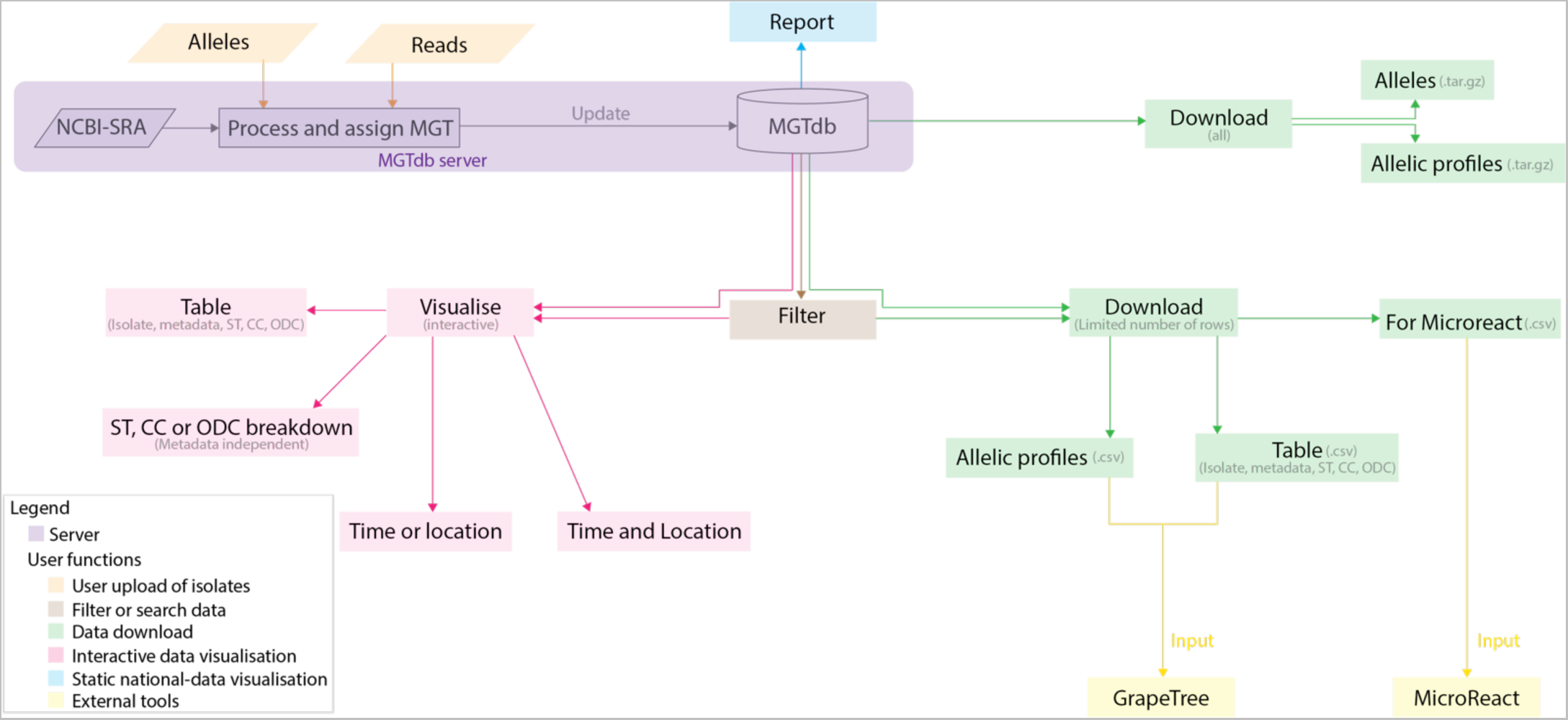
Data visualisations and downloads in MGTdb. The MGTdb database consists of isolates - user-uploaded or those retrieved from NCBI-SRA - assigned the MGT identifiers. The MGTdb web-service contains features to filter (indicated by brown), visualise (pink and blue) or download (green) these isolates. Indicated by pink: isolates are displayed in a table, either as a complete set or filtered with by metadata, ST, CC, and ODC assignments. This data can also be explored via three interactive graphs. The first enables exploring the set of isolates and their MGT assignments in the context of either time or location, the second, in the context of both, and the third interactive graph does not use metadata, and enables exploration of isolates and their MGT assignments. Indicated by blue: a report of country- wise or project-wise static summaries of isolates for the last 10 years, at each MGT level, can be generated. Indicated by green: the first 2000 isolates in the displayed table can be exported for use with Microreact - a web-tool which enables phylogeographical and temporal exploration of isolates. The data displayed in the initial table, and the allelic profiles for the largest MGT-level STs can also be downloaded. For these two files, the current limits are set to 1,000,000 rows (essentially the entire database) for the tabular data, and 10,000 for allelic profiles. The complete set of publicly available isolates, their allelic profiles and allele sequences can be downloaded separately from the organism main page.

Interactive graphs available on MGTdb enable dynamic exploration of isolates and their MGT associated assignments. As the information made available on MGTdb is very dense, three types of interactive visualisations are currently made available. The first summarizes isolate counts for STs (or CCs, or ODCs) at any particular MGT level by either time (**Figure 2a**) or location (**Supplementary Figure S3)** as a stacked bar chart. The second visualisation summarizes counts of isolates, counts of STs (or CCs, or ODCs) at any particular MGT level, or both (using the Shannon diversity index(26)), in both time (x-axis) and location (y-axis), as a heatmap (**Figure 2b**). The third interactive graph summarizes isolate counts over STs (or CCs, or ODCs) at all MGT levels simultaneously (**Supplementary Figure S4**). It does not utilize any additional metadata, and only utilizes the MGT associated assignments. In all these graphs, the interactivity enables easy switching of displayed data, to visualize isolate counts by the MGT levels, STs, CCs or ODCs, and the locational and temporal breakdowns - thus enabling easy exploration of isolate relationships at a wide variety of resolutions and contexts.

**Figure 2.**
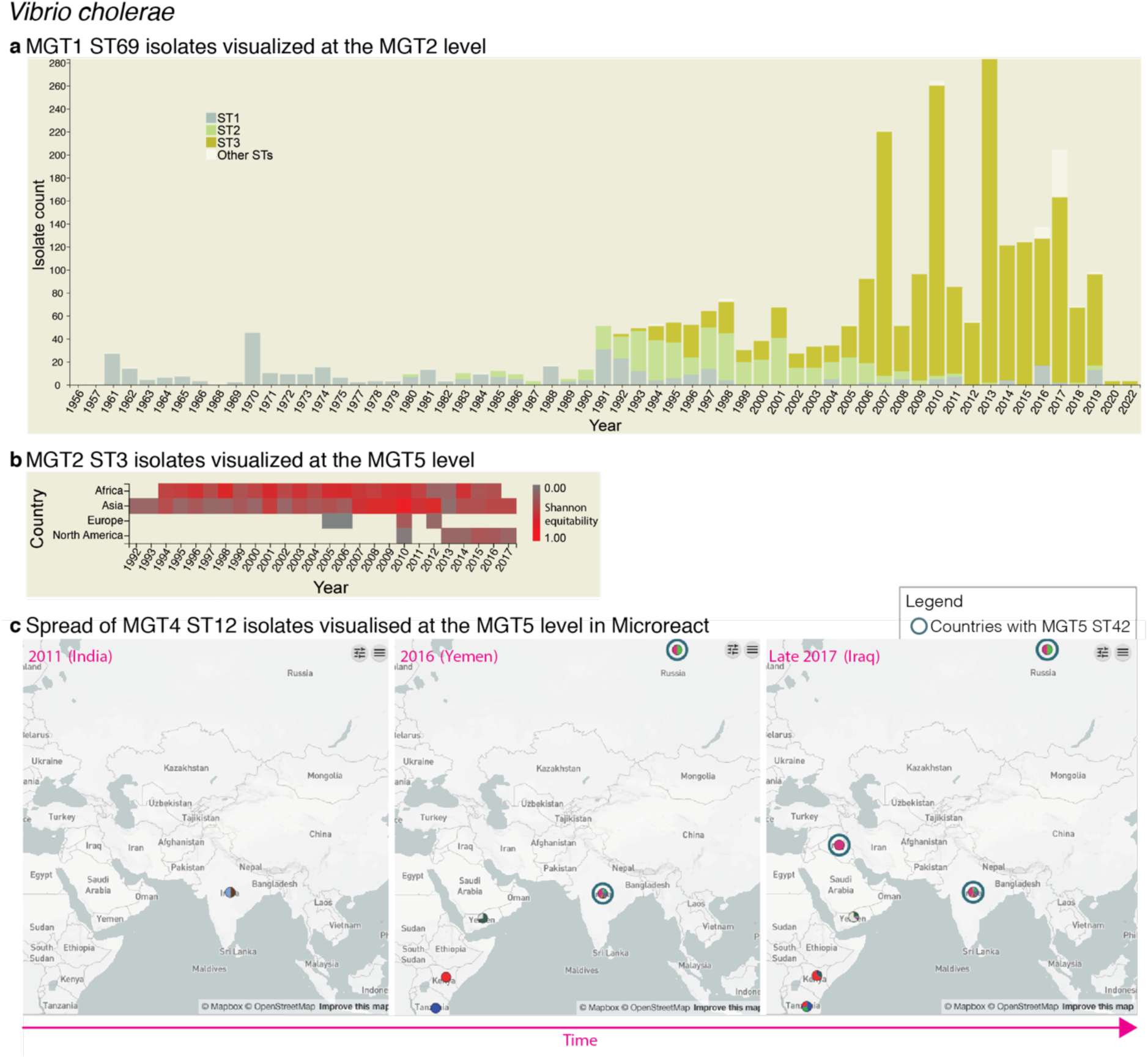
Interactive charts and streamlined integration with Microreact enabled in MGTdb. **a**. An interactive chart in MGTdb that enables summarising all or filtered isolates at any MGT level using either the locational or the temporal metadata. The chart shows the current cholera pandemic isolates (i.e. MGT1 ST69 or seven-gene MLST ST69) at the MGT2 level. This data can be explored on MGTdb at https://mgtdb.unsw.edu.au/vibrio/isolate-list?mgt1=69&searchType=and (data view level MGT2 ST, with ‘Only highlight top’ set to ‘3’). **b**. An interactive chart in MGTdb that enables summarising all initially-loaded or filtered isolates using both locational and temporal metadata. The chart shows the current cholera wave isolates (i.e. MGT2 ST3) at the MGT5 level. This data can be explored on MGTdb at https://mgtdb.unsw.edu.au/vibrio/isolate-list?ap2_0_st=3&searchType=and (data view level MGT5 ST). **c**. Microreact visualisation of MGT4 ST12 isolates exported from MGTdb. A data file - wherein the latitude and longitude are appended, along with unique colours for STs - can be exported from MGTdb for use with Microreact. Microreact enables visualisation of isolates at any chosen MGT level, both geographically and temporally. Shown here is the temporal step-through (as enabled on Microreact) of MGT4 ST12 isolates, visualized at the MGT5 level. The data shown in this figure can be explored in Microreact using the link https://microreact.org/project/bjdgCuWdVowtJHwq2npYxi-mgt5-st12-vibrio-cholerae and selecting ‘Colour Column’ as MGT5. Alternatively, the original data can be viewed in MGTdb using the link https://mgtdb.unsw.edu.au/vibrio/isolate-list?ap4_0_st=12&searchType=and.

For national surveillance, a report can be easily generated depicting trends of the most frequently observed STs within a nation. The report contains static visualisations systematically summarising isolates at all MGT levels for the selected country within a selected range of years (**Figure 3**). Each time a report is generated, the latest data, comprising all available public isolates upto the moment of request, are retrieved from the database. Thus, a snapshot of the trends observed at a particular time can be saved by downloading the generated report.

**Figure 3.**
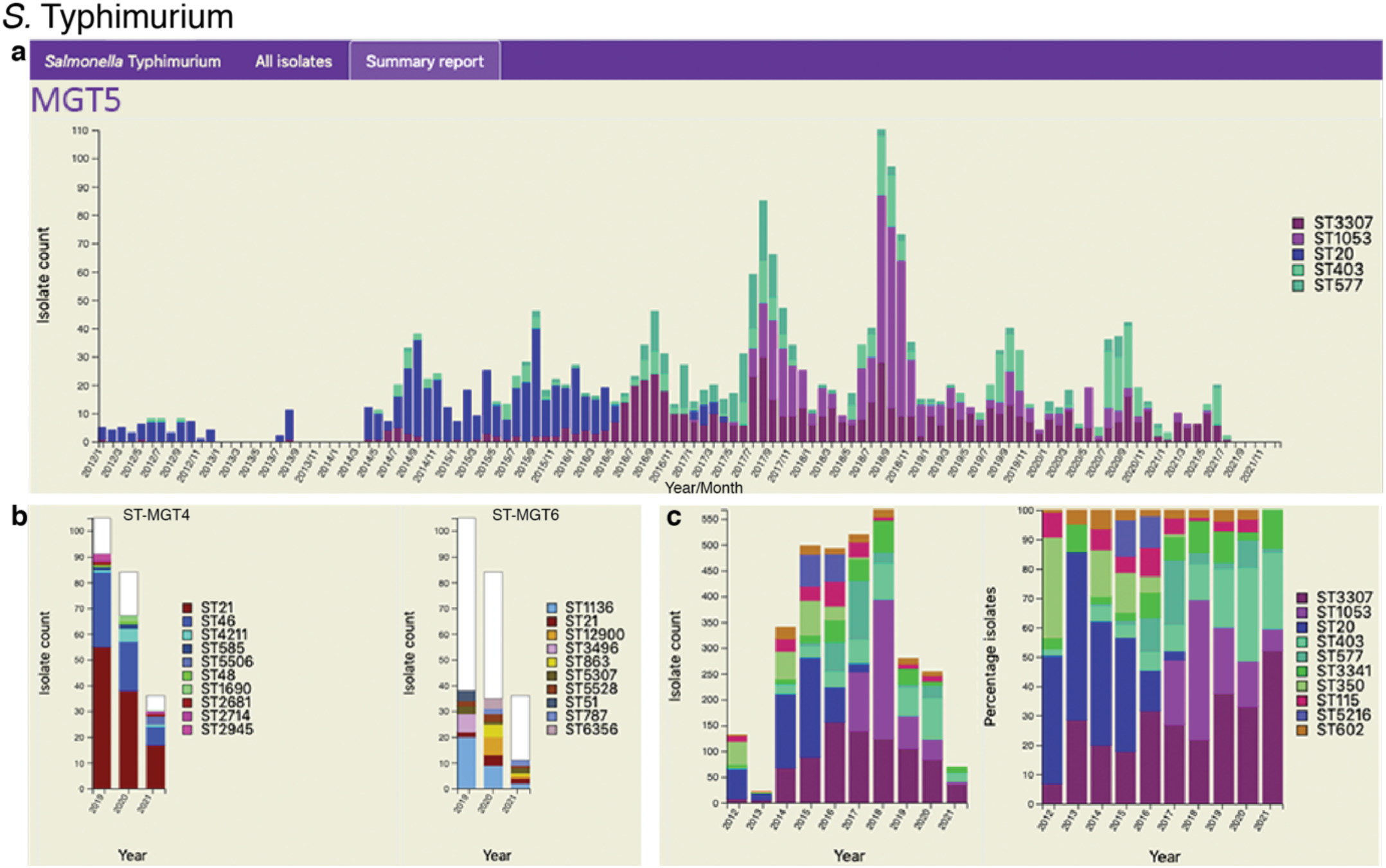
Static charts from the report feature on MGTdb which enables national surveillance. For any country, or any user’s project (which houses a set of isolates), a report can be generated. The report presents isolates (and MGT assignments), within a specified country or project for a specified range of years, as general summaries, as well as, summarizes the isolates at each individual MGT level as a series of graphs. Shown here are graphs from the report generated for *S*. Typhimurium MGT5 STs in the UK over the past 10 years (2012–2021). In particular, **a**. summary of top five MGT5 STs by month, **b**. summary of MGT5 ST3307 isolates at the MGT4 and MGT6 levels, and **c**. summary of top ten MGT5 STs by year, where the first graph shows isolate counts, and in the second, the counts have been scaled by the maximal ST distribution observed that year. This data can be explored on MGTdb at https://mgtdb.unsw.edu.au/salmonella/summaryReport by choosing the country ‘United Kingdom’, setting the ‘Year start’ as 2012, ‘Year end’ as 2021, and then generating the report.

MGTdb enables download of the displayed data (e.g. as displayed in **Figure 4a**). In particular, all or filtered isolates and their metadata, along with their MGT associated assignments can be downloaded. Additionally, allelic profiles of the largest scheme can be downloaded which, in conjunction with the previously mentioned file, can be used to generate neighbour-joining or minimal-spanning trees, produced by tools such as GrapeTree (27) - as shown in **Figure 4b**. Such a tree can then be coloured according to the assigned STs, CCs or ODCs at any MGT level, or any other metadata. MGTdb also enables download of a file for import into Microreact (28) - a tool which enables visualising the phenotypic, epidemiological, temporal and geographical distribution of isolates. For import into Microreact, MGTdb appends the latitude and longitude for a country, and unique colours of STs, for visualisation. The resulting file can then be effectively explored in Microreact as exemplified in **Figure 2c**. Lastly, all allele sequences for a species, and allelic profiles of public isolates, are collected, compressed and made available for each hosted organism daily (at https://mgtdb.unsw.edu.au/salmonella/, https://mgtdb.unsw.edu.au/enteritidis/, and https://mgtdb.unsw.edu.au/vibrio/).

**Figure 4.**
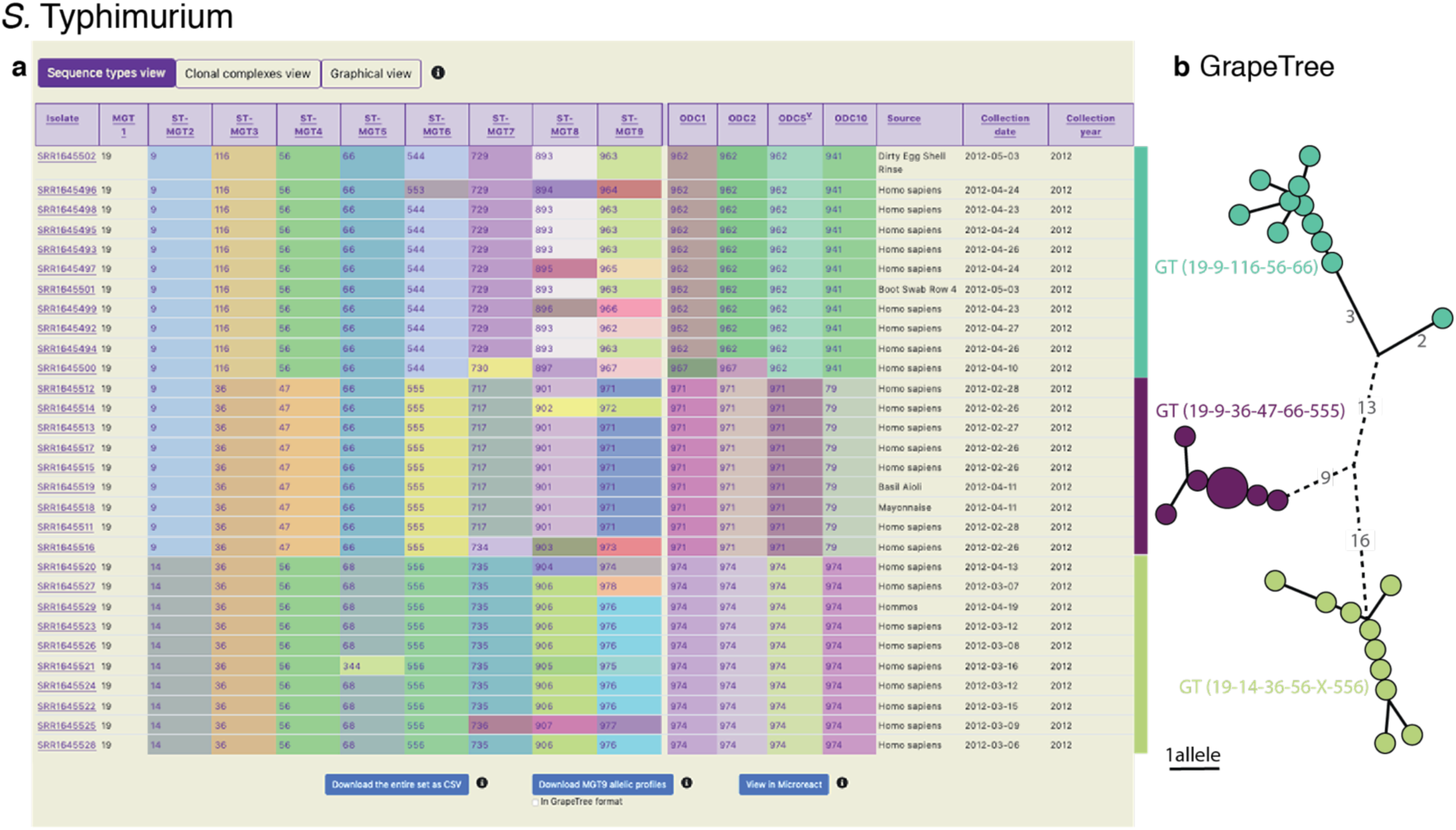
Display of all data in MGTdb and exported data enabling streamlined integration with GrapeTree. **a**. All isolates - initially-loaded or filtered - are shown in an interactive table. Shown in the table are also the sequence types (STs) assigned to the isolates at all MGT levels and outbreak detection clusters (ODCs). This view can be switched to visualise clonal complexes (CCs), instead of STs, for all MGT levels, using buttons above the table. The CCs group STs by single linkage clustering with a 1 allele difference cutoff, and ODCs group the largest MGT-level STs, with 1, 2, 5 and 10 allele difference cutoffs. The table shows a maximum of 100 isolates per page. To summarise all the initially-loaded or filtered data, a user can switch to graphical view (button displayed above the table), or download the data (using buttons below the table). The user can download the data displayed in the table (MGT associated assignments), allelic profiles for the largest MGT level, or export the data to Microreact. **b**. GrapeTree visualisation of isolates exported from MGTdb. The allelic profiles and the MGT assignments can be imported into GrapeTree to build and visualise a neighbour joining graph (as shown here; or a minimal spanning tree). The data shown in the table and the GrapeTree network graph are from three outbreaks of *S*. Typhimurium that occurred in NSW, Australia in 2012, and can be explored on MGTdb at: https://mgtdb.unsw.edu.au/salmonella/isolate-list?year=2012&state=NSW&searchType=and.

## 3. Case studies

In this section, we demonstrate the utility of MGTdb to enable epidemiological studies at global and local scales, as well as over long- and short-timespans, via three case studies. In the first study, the seventh pandemic isolates of *V. cholerae* are examined, with a particular focus on the STs associated with the 2016 to 2017 cholera outbreak in Yemen (**Section 3.1**). In the second study, the distribution of *S*. Typhimurium isolates in UK is examined (**Section 3.2**). Finally, in the third case study, outbreaks caused by *S*. Typhimurium in Sydney, Australia, are examined (**Section 3.3**).

### 3.1 The global population structure of *V. cholerae*

The population structure of *V. cholerae* has been described using MGT (24). The ongoing seventh pandemic of cholera is predominantly caused by the seven-gene MLST ST69 (also referred to as MGT1 ST69). At the MGT2 level, the seventh pandemic isolates are divided into three predominant STs, namely MGT2 ST1, ST2, and ST3, representing the three distinct waves of the pandemic. Interactive graphics available in MGTdb were used to visualize such trends - **Figure 2a** shows the temporal distribution of these three MGT2 STs. The temporal and geographical distribution of the most recent wave of isolates, MGT2 ST3, are shown in **Figure 2b**. As the latter figure shows, the current wave originated from Asia from where it then spread to Africa, Europe and America; high ST diversity (at the MGT5 level) is observed in Asia and Africa.

MGTdb enables exporting data for use with Microreact (28) - a web-based tool that enables visualisation of geographical epidemiology of isolates by overlaying ST distributions on a map, along with temporal epidemiology by enabling a temporal walk-through. The exported data from MGTdb can be used to study isolates at any level of resolution within Microreact. As an example, consider the MGT4 ST12 isolates - the ST associated with the 2016 to 2017 cholera outbreak in Yemen (24) - shown in **Figure 2c** at the MGT5 level. Weill *et al*. (29) studied the genomic epidemiology of these outbreak isolates. They found that these isolates were phylogenetically distinct from isolates observed in Iraq (2007 to 2015) and Iran (2012 to 2015), and closer to the 2015 to 2016 isolates from Kenya, Tanzania and Uganda. Indeed, observing the temporal global dissemination of the MGT4 ST12 isolates (**Figure 2c**), this ST is not found in Iran, and in Iraq the MGT4 ST12 isolates are observed in late 2017 - after the start of the Yemen outbreak. Furthermore, MGT5 ST42 isolates, indicated by blue circles, observed in Iraq are observed in India and Russia, but not Yemen, suggesting that this ST may have been imported from India/Russia rather than Yemen. Thus, by exploring data from MGTdb in Microreact, users can easily combine the unique features of MGT and Microreact for studying the global temporal epidemiology.

### 3.2 *S*. Typhimurium in UK

The foodborne pathogen *S*. Typhimurium has been described using MGT (21). Isolates described via MGT can be studied at different resolutions for temporal and geographic trends. For national-level surveillance, reports for any chosen country can be generated in MGTdb, consisting of static charts summarising isolates at each MGT level within a selected range of years (**Section 2.7**) - a user can then easily identify the best level for examining relevant epidemiological patterns. As an example of long-term national surveillance, **Figure 3** summarises isolates from UK in the last 10 years (2012-2021) at the MGT5 level. The presentation of *S*. Typhimurium distribution by month (**Figure 3a**) highlights isolates with MGT5 STs that have become extinct (e.g., ST20), STs that are newly emerging (i.e. ST3307 and ST1053) and STs that are persistently present (ST403 and ST577). **Figure 3b** shows how STs are distributed at the previous and next MGT level for MGT5 ST3307 isolates (the most common MGT5 ST in the UK over the past 10 years). **Figure 3c** depicts the top 10 STs by year, where the first graph shows distribution of isolates, and the second shows distribution of isolates scaled to 100%. Here, the former graph indicates sampling rates, and the latter reveals trends in ST frequency over time.

### 3.3 *S*. Typhimurium outbreaks in Sydney, Australia

Lastly, we demonstrate MGTdb’s utility for examining short-term epidemiology and outbreaks using *S*. Typhimurium. Octavia *et al*. (30) studied the genomic epidemiology of *S*. Typhimurium isolates that caused five outbreaks in NSW, Australia. Of these, three outbreaks occurred in 2012, namely outbreaks *2* (for which 11 strains were sequenced), *4* (9 strains), and *5* (10 strains) shown in the table in **Figure 4a**. These isolates were identified by using MGTdb’s search functionality (**Section 2.5**) to filter the database (which contains all publicly available isolates): setting ‘Year’ to ‘2012’ and ‘State’ to ‘NSW’. These three outbreaks can be uniquely and stably characterised via MGT as the following genome types (by observing the table shown in **Figure 2a**), GT19-9-116-56-66 (i.e., MGT1 ST19, MGT2 ST9, MGT3 ST116, MGT4 ST56 and MGT5 ST66), GT19-9-36-47-66-555 and GT19-14-36-56-X-556, respectively.

Octavia *et al*. also found that *S*. Typhimurium genomes differed by a maximum of five single nucleotide polymorphisms (SNPs) for outbreak 2 and by a maximum of 1 SNP for outbreaks 4 and 5. This relationship can be simply described by outbreak detection clusters (ODCs) - ODC5-962 groups outbreak 2 isolates together, while ODC1-971 and ODC1-974 groups isolates in outbreaks 4 and 5 together respectively. To visualize the relationships revealed by single linkage clustering methods, such as ODCs, MGTdb enables the download of allelic profiles for the largest MGT level (e.g. MGT9 for *S*. Typhimurium). These downloaded allelic profiles can be used to generate a neighbour joining or a minimal spanning tree using tools such as GrapeTree (27), and the downloaded MGT associated assignments (i.e. the table seen in **Figure 4a;** downloaded by clicking on ‘Download entire set as CSV’) can be used to visualise the relationships within the context of the MGT assignments (**Section 2.7**). For example, the neighbour joining tree of these outbreak isolates - shown in **Figure 4b** (coloured by ODC5) - indicates the same relationship between isolates as revealed by Octavia *et al*. (albeit using allele differences rather than SNP differences).

## 4. Summary and conclusions

In this paper, a web-service, MGTdb, is presented. MGTdb enables a user to upload sequenced reads or locally extracted alleles of isolates, obtain MGT assignments and examine the isolates in the context of all isolates in the database. Additionally, MGTdb regularly retrieves and processes publicly available genome sequencing data from NCBI-SRA (25) - thus enabling the study of essentially all publicly available isolates for a species. Visualisation tools are presented to assist analysis at a wide variety of resolutions, such as long-term or short-term, along with capabilities to download a wide variety of data - thus enabling downstream and alternative types of epidemiological studies.

Key characteristics for building a usable and successful nomenclature system have been previously identified [82, 161, 162]. MGTdb has been developed in line with these recommendations. In particular, the key characteristics as implemented in MGTdb are outlined below:

1. Comprehensive characterization of target bacterial populations.
2. Standardised and stable nomenclature for the characterisation of isolates.
3. Standardised and stable nomenclature for the characterisation of relationships between isolates.
4. Database system for data management.
5. Enables data mining and analysis.
6. Sufficient computational resources for nomenclature assignment, data storage and management, and data analysis.
7. The nomenclature, and analyses can be downloaded.
8. Open source, open access, and free of cost.
9. User login to track uploaders and preserve privacy when required.
10. Enables uploading raw reads or locally extracted alleles. The upload of alleles enables further privacy preservation, as only the sequence needed for typing is submitted. Lastly, the uploaded data is quality controlled.

Currently, one major limitation of MGTdb, as recommended for a nomenclature system, is to enable obtaining assignments remotely (i.e. without the need to submit the isolates), such as implemented in chewie-NS [163]. Approaches for enabling this feature in MGTdb are currently being explored.

In conclusion, MGT presents a powerful approach enabling researchers to investigate sequenced bacterial isolates at multiple levels of typing resolution, in the local and global context. Through this method, epidemiological investigations such as those determining if isolates belong to previous, ongoing or new outbreaks, epidemics or pandemics can be easily performed. To this end, the MGTdb web service enables a user to submit isolates and obtain MGT associated assignments. It contains intuitive and easy-to-use capabilities for visualisation, file export and download. MGTdb is currently available and ready to use for three exemplar organisms - with schemes for more species in development. A locally installable version of the database is available for further development and hosting of MGT schemes and for users to develop their own MGT schemes of any pathogens.

## Acknowledgements

The authors would like to thank Robin Heron for providing fantastic support with Katana and the virtual machines. The authors also thank Liam Cheney and Laurence Luu for helpful discussions. The authors are also grateful to members of the Health Protection NSW, especially Jennie Musto, Keira Glasgow and Daneeta Hennessy, and the Institute of Clinical Pathology and Medical Research, and, Qinning Wang and Peter Howard from NSW Health Pathology for testing and providing valuable feedback regarding various aspects of usability of MGTdb. Finally, the authors acknowledge the NHMRC grants (grant numbers 1129713 and 211806) that supported this work.

## Conflicts of interest

None

## Notes

https://mgtdb.unsw.edu.au

### Competing Interest Statement

The authors have declared no competing interest.

### Summary of Updates

Uploaded document as PDF to improve image quality.

