## Supplementary for "MGTdb: A web service and database for studying the global and local genomic epidemiology of bacterial pathogens"

#### Supplementary figures

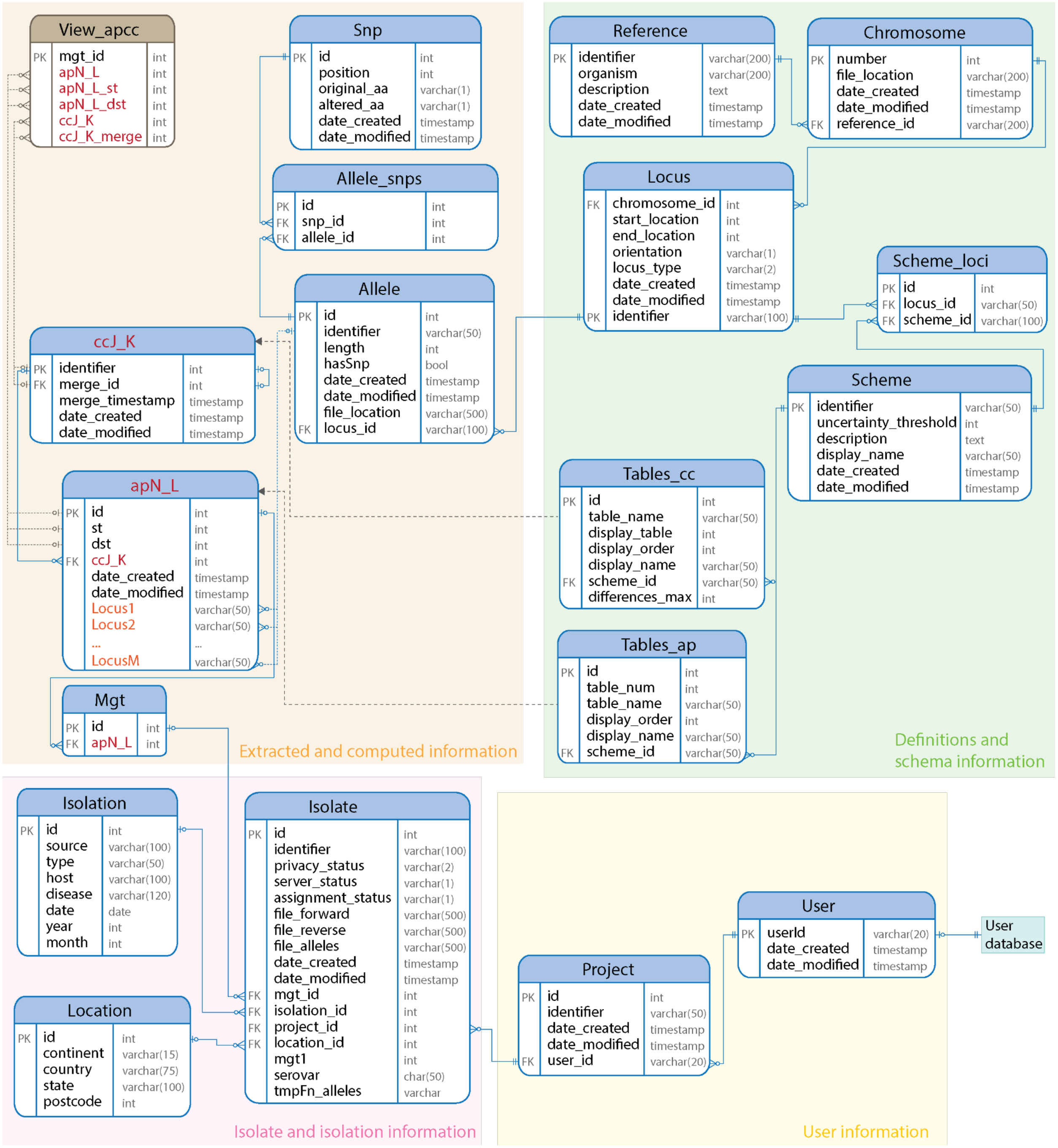


**Supplementary Figure S1.** Organism database schema. Data for each organism is stored in an organism-specific database on the MGTdb web server. Four types of information are stored: 1) *user,* 2) *isolate and isolate metadata,* 3) *definitions and schema* and 4) *extracted and computed,* shown here using different background colors. The *definitions and schema* tables store general information regarding the organism, as well as the loci and the MGT scheme definitions. For each MGT-level scheme, a table (table indicated as `apN_L`, in *extracted and computed information*) is created to store the allelic profile and sequence type assignment, at the MGT level, for an isolate. Data pertaining to these tables is contained in `Tables_ap`. For each scheme, a variable number of clonal complexes can be computed. The default is one clonal complex computed for each scheme, allowing a maximum of 1 allele difference, with the exception of the largest scheme, for which four complexes are computed at maximum allowed differences of 1, 2, 5 and 10 alleles. Data for each clonal complex is added to its own table (indicated as `ccJ_K`), and data pertaining to these tables is contained in `Tables_cc.` Thus, organism-specific tables and column names are indicated by red and orange in this database schema. *User information* tables store information regarding any projects that a user creates. Any number of isolates can be created within a project. Isolates, and any associated metadata, such as temporal and locational information are stored as part of the *isolate and metadata* tables. For each isolate, it's assigned MGT, STs (along with allelic profiles, alleles and allele variants, such as SNPs) and CCs for each MGT level are stored as part of the *extracted and computed information* tables.

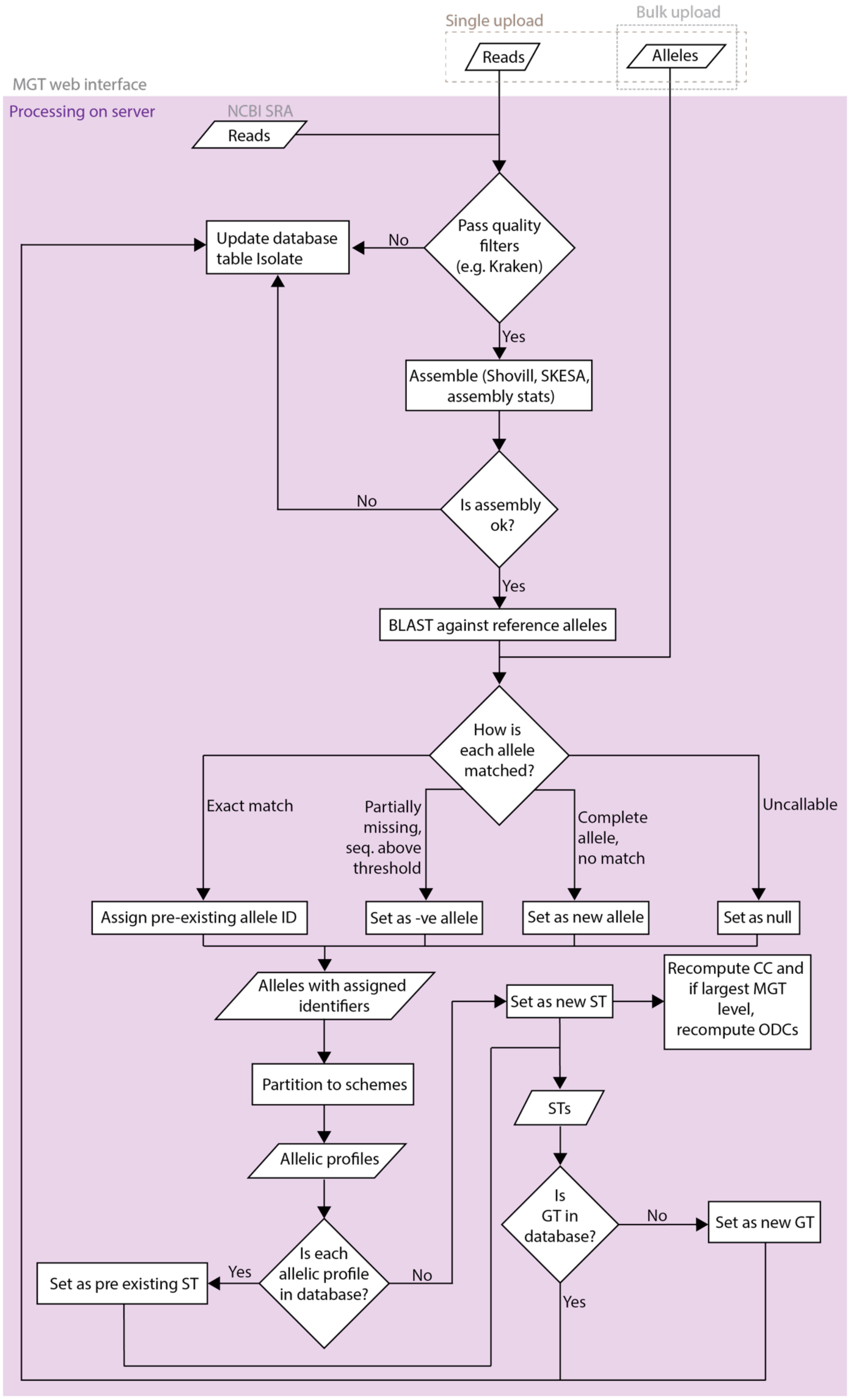

Single upload
Bulk upload
Reads
Alleles
MGT web interface
Processing on server
NCBI SRA
Reads
Update database table Isolate
No
Pass quality filters (e.g. Kraken)
Yes
Assemble (Shovill, SKESA, assembly stats)
No
Is assembly ok?
Yes
BLAST against reference alleles
How is each allele matched?
Exact match
Partially missing, seq. above threshold
Complete allele, no match
Uncallable
Assign pre-existing allele ID
Set as -ve allele
Set as new allele
Set as null
Alleles with assigned identifiers
Set as new ST
Recompute CC and if largest MGT level, recompute ODCs
Partition to schemes
STs
Allelic profiles
Is GT in database?
No
Set as new GT
Set as pre existing ST
Yes
Is each allelic profile in database?
No
Yes

**Supplementary Figure S2.** Isolate processing and assignment of MGT associated identifiers. Inputs and key steps involved in data processing on the MGTdb web server. Once logged in, a user can upload sequenced reads, or alleles - where the alleles have been extracted using the MGT pipeline[1], locally, by the user. Due to smaller file size, the latter option allows a user to upload multiple isolates at once. Additionally, newly deposited Illumina sequenced reads are also periodically retrieved from NCBI. If reads are uploaded by the user, or, for reads retrieved from NCBI, alleles are extracted on the compute cluster associated with the web-server. The uploaded or extracted alleles are then compared with alleles in the organism-specific database. Any new alleles identified are added to the database, along with the resulting allelic profile and sequence type, and an MGT assignment to the isolate is made. For each MGT level where new alleles are added, clonal complexes (and outbreak detection clusters) are recomputed. The isolate with its assigned MGT STs, CCs and ODCs can then be searched and visualised in the context of the global and the local database. A detailed presentation of the key steps involved in data processing can be found at https://mgt-docs.readthedocs.io/en/latest/analysis_pipeline.html.

[1] MGT pipeline for extracting alleles from reads: https://github.com/LanLab/MGT_reads2alleles (last visited on 2021, September 13).

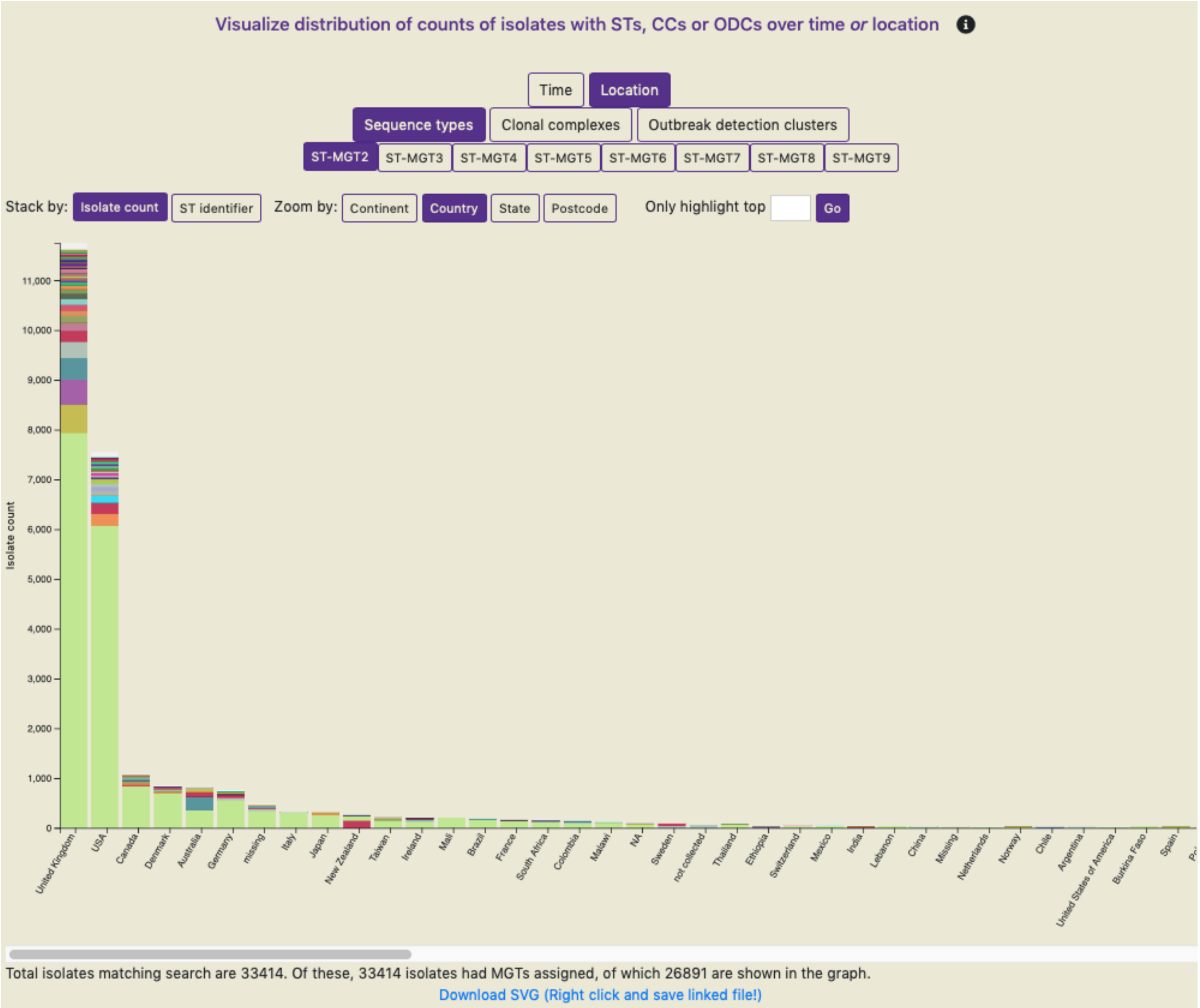


**Supplementary Figure S3**. Screenshot of an interactive graph enabled on MGTdb in the 'Graphical view' tab (see **Figure 4a** for location of Graphical view). This graph summarizes isolate counts over STs (or CCs, or ODCs) at a particular MGT-level on either time or location (as shown here; countries are sorted from left to right in descending order of isolate counts) as a stacked bar chart. The interactivity, enabled by the various buttons (shown at the top), enables a user to change how the data is plotted. For example, a user can choose to visualise data at the continent, country, state (sub-country) or postcode level. Furthermore, a user can choose to change the MGT level, the MGT assignment type (ST, CC or ODC), change how the bars are stacked (by count, or by identifier) or only highlight bars with a user-specified number of top ST distributions. Through the interactive switching, data is transformed and replotted - not all isolates visualized may contain the specified metadata for plotting. Hence, the number of isolates annotated with the required metadata and shown in the graph are indicated below the plot. Lastly, the displayed plot can be saved to a local computer as a scalar vector graphics (SVG) file, and edited in tools such as Adobe Illustrator. The data shown in this figure are all the publicly available isolates of *S.* Typhimurium.

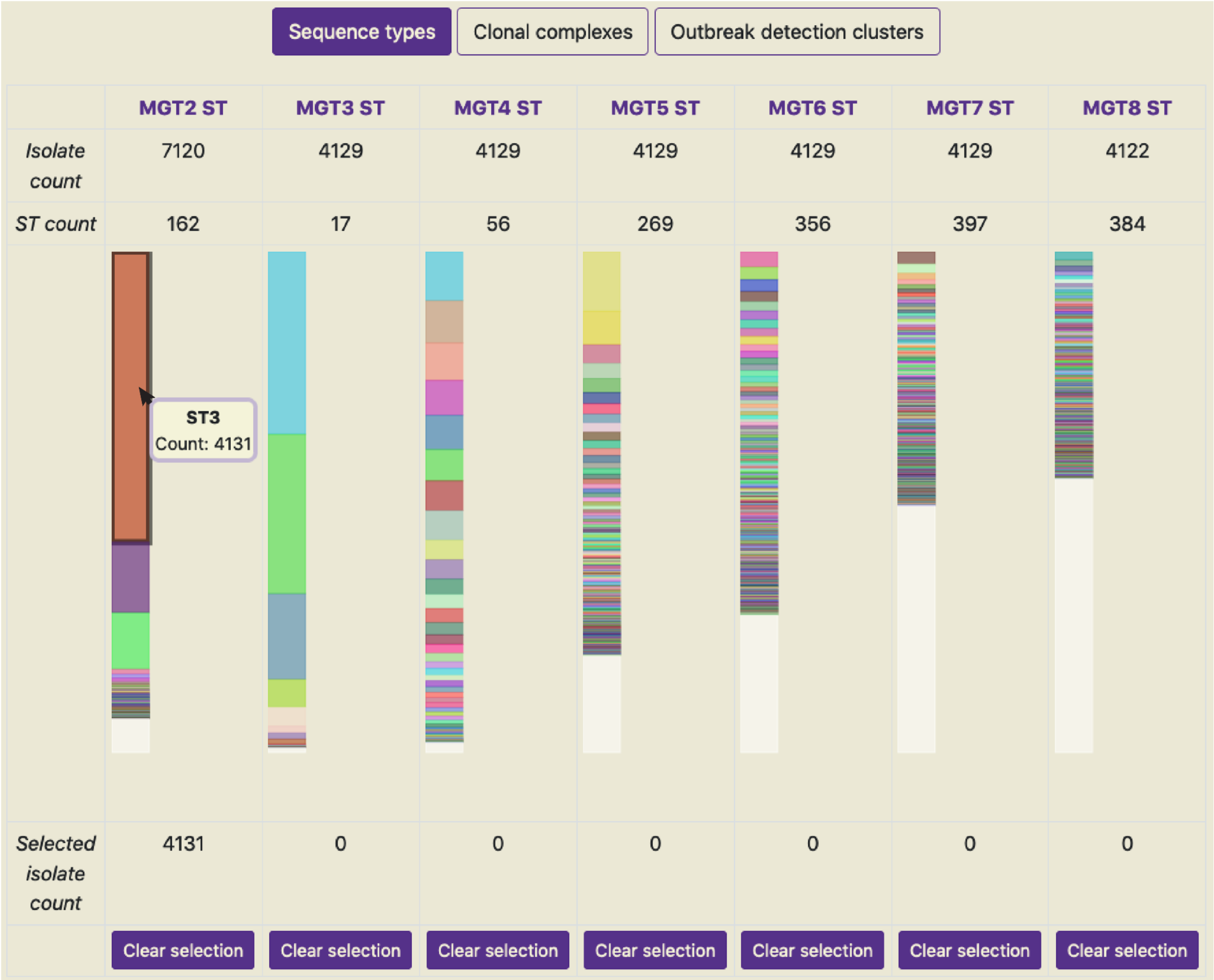


**Supplementary Figure S4**. Screenshot of an interactive graph enabled on MGTdb in the ‘Graphical View’ tab. This graph summarizes isolate counts over STs (or CCs, or ODCs) at all MGT-levels. For each MGT level, the first row indicates isolate counts and the second row indicates ST counts. The third row shows a stacked bar chart depicting ST distributions of isolates. Hovering over any bar, in the stacked bar chart, reveals the ST identifier and the number of isolates (as seen here for MGT2 ST3). Any bar can be clicked, which selects the ST value - this results in the replotting of the stacked bars at all other MGT levels such that the ST distributions are plotted for those isolates which are assigned the selected ST at the selected MGT level. Additionally, the number of isolates assigned to the selected ST are shown in the fourth row. Bars can be selected at any MGT level, and in any order, with the exception of the white bar in each level. The white bars indicate distributions of STs assigned to single isolates. These selections can be cleared by re-clicking on the bars, or using the button ‘Clear selection’. The data shown in this figure are the publicly available isolates of *V. cholerae*, and MGT2 ST3 has been selected.

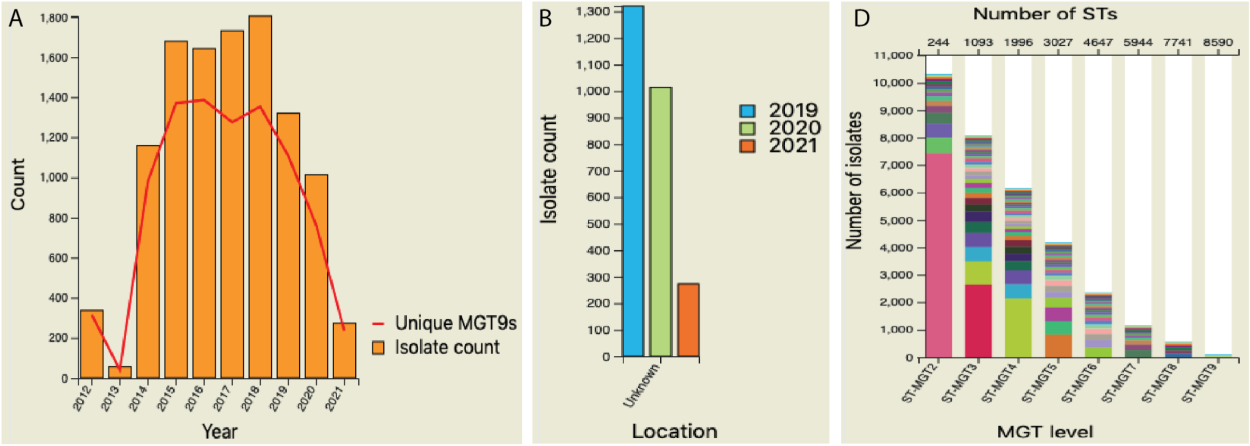


C

| Year | Number of isolates | Number of ST-MGT2s | Number of ST-MGT3s | Number of ST-MGT4s | Number of ST-MGT5s | Number of ST-MGT6s | Number of ST-MGT7s | Number of ST-MGT8s | Number of ST-MGT9s |
|---|---|---|---|---|---|---|---|---|---|
| 2019 | 1320 | 78(31) | 290(134) | 435(239) | 600(366) | 791(602) | 891(753) | 1028(977) | 1106(1064) |
| 2020 | 1014 | 51(14) | 184(68) | 276(131) | 373(207) | 489(340) | 574(462) | 709(641) | 760(718) |
| 2021 | 273 | 29(4) | 78(14) | 112(44) | 135(68) | 189(115) | 202(160) | 223(188) | 234(212) |

**Supplementary Figure S5.** Screenshots of four charts, giving an overview of isolates for a chosen country or project, contained in reports generated on MGTdb. For any chosen country or project, and a specified year range, four overview charts are systematically generated: A) depicts isolate counts per year and the number of unique largest-level MGT STs; B) depicts counts by sub-country or state (if the report is generated for a country), or breaks down isolates by country (if the report is generated for a project); C) summarizes for all MGT levels - within the past three years - the number of isolates, the number of STs and the number of STs newly observed in a particular year when compared to STs in the entire retrieved dataset; D) depicts ST distributions for isolates at each MGT level. The report was generated here for the organism *S.* Typhimurium, the country UK and year range set to 2012 to 2021. The figure was then generated by taking screenshots of the report, which were combined in Adobe Illustrator.

### Supplementary notes

#### S1 Data visualization and download

The complete list of data visualization techniques, and downloads enabled in MGTdb, are summarized in **Figure 1**. Here, we describe these in detail.

On the MGTdb web-server, any initially-loaded or filtered isolates, along with their MGT assignments (namely STs, CCs, and ODCs) are shown in an interactive table (see **Figure 4a**). Key features of the table are described here.

- A quick search can be launched by clicking on any ST, CC or ODC identifier in the table.
- By default, each cell containing a ST, CC, or ODC identifier is uniquely colored based on the value of the identifier. This enables easy visualization and recognition of isolates with similar

STs, CCs or ODCs across the table, as compared to using the numeric identifiers for this purpose. This setting can be easily switched off when required.

- Certain allelic profiles may contain alleles which pass all quality filters, but may still contain a small proportion of missing or indeterminable sequence. Such alleles are called negative alleles, and the resulting STs are called degenerate STs (dSTs). STs containing dSTs are designated as ST.dST; an example is MGT5 ST4.11 (https://mgtdb.unsw.edu.au/salmonella/isolate-list?ap5_0_st=4&ap5_0_dst=11). Here, the additional dST indicates that, in comparison to MGT5 ST4, there is no difference in the sequences for the two allelic profiles, apart from missing information in one or more loci in the allelic profile assigned dST. A dST designation is intended to alert the user that there is some degree of missing information involved in the ST assignment. By default the dSTs are hidden, but can be switched on when required.
- Finally, the list of initially-loaded or filtered isolates can be sorted by any column, enabled by clicking the table header. Here, rows containing null values for the sorted column are added to the end.

Currently, the table is set to show a maximum of 100 isolates per page. To summarize all the initially-loaded or filtered data, a user can switch to graphical view, or download the data. As shown in **Figure 4a**, these buttons are found above or below the table, respectively.

A user can download the data displayed in the table (described above), allelic profiles for the largest MGT level, or export the data for Microreact (1). During export for Microreact, a tabular file is downloaded, which can be manually uploaded to Microreact by the user; this option enables the user to modify the file as required before visualizing the data in Microreact. MGTdb appends the geolocational information, i.e. the latitude and longitude for a country, as well as unique colour values for STs. An example of data exported from MGTdb and visualized in Microreact can be seen in **Figure 2c**. The allelic profiles downloaded from MGTdb can be used to generate a minimal spanning tree (MST) in tools such as GrapeTree (2) (**Figure 4b**). By default the downloaded allelic profiles file contains negative alleles. However, for the purposes of building a MST, the user may want to ignore the missing information in alleles (which is necessary for Grapetree). Hence, MGTdb allows a user to download the allelic profiles where the negative alleles are shown as their complete counterpart (i.e. are treated as a full-called allele) - by selecting the option ‘In GrapeTree format’ below the ‘Download MGT9 allelic profiles’ button, as seen in **Figure 4a** below the table. The MGT-associated-assignments tabular file can also be input into GrapeTree, where the assignments can serve as the metadata for colouring the graph (for example, according to a particular MGT level - thus enabling the visualization of ST distribution in the tree). Each of these three types of downloadable-files have limits in the number of rows that can be downloaded (to enable downloads within a reasonable time). Currently, these limits are set to a maximum of 1,000,000 rows for the MGT associated assignments file (this is currently effectively the complete data as counts of isolates of all hosted organisms is below this limit), a maximum of 10,000 rows for the allelic profiles file, and 2,000 rows for the Microreact file. To circumvent this limitation, a user can download the allelic profiles for the largest MGT level (as the schemes have currently been set up such that the largest MGT level includes all loci from other MGT levels) from the main organism page, or allelic profiles from the project page (these contain the project specific allelic profiles, as any private data is not available on the organism main page). Additionally, a user can also download all the allele sequences from the organism main page. The allele sequences can be used for various downstream analysis such as SNP analysis.

Apart from downloading the data, a user can also visualize the initially-loaded or filtered isolates and MGT associated assignments in interactive graphs on MGTdb. As the information made available on MGTdb is very dense, three types of interactive visualisations are currently made available.

The first summarizes isolate counts over STs (or CCs, or ODCs) at a particular MGT-level on either time (e.g. **Figure 2a**) or location (**Supplementary Figure S3**) as a stacked bar chart. The interactivity enables a user to switch the time or location zoom level (for example, a user can zoom-in to view the distributions from year, to per month, or per day), change the MGT level, the MGT assignment type (ST, CC or ODC), change how the bars are stacked (by count, or by identifier) and many more. **Supplementary Figure S3** depicts the various buttons which enable this interactivity.

The second visualisation summarizes counts of isolates, counts of STs (or CCs, or ODCs), or both, in both time (x-axis) and location (y-axis), as a heatmap. Summarizing the counts of isolates, or STs (or CCs, or ODCs) is straightforward in the heatmap. To summarize both isolate and ST counts, the Shannon equitability index ($E_H$) (3) is used (example **Figure 2b**). It is calculated in MGTdb as follows:
For a given time and location, let *S* be the total number of STs (or CCs, or ODCs); let *i* be an ST value, $c_i$ be the number of isolates assigned $S_i$ and *T* be the total number of isolates then the proportion of isolates assigned a particular ST ($p_i$) is calculated as $p_i = \frac{c_i}{T}$; and the Shannon diversity index (3) as $H = -\sum_{i=1}^{S} p_i \times ln\,(p_i)$; and finally the Shannon equitability ($E_H$) as $E_H = \frac{H}{H_{max}}$; where $H_{max}$ is the maximum H calculated over the entire visualised dataset (i.e. across all time and location pairs visualized). The $E_H$ value lies between 0 and 1, for lowest to highest evenness. For example, 1 indicates high isolate count and high number of STs.

Similar to the first interactive visualization, the second visualisation also enables interactive switching and exploration of data. The data displayed in both the first and second interactive graphs depends on metadata annotations by the users. MGTdb requires that a user provides minimum metadata of country and year. However, genome data retrieved from genome databases such as NCBI may not contain these metadata. Hence, it is worth noting that effective data exploration will be dependent on the metadata annotation available for the isolates being studied by the user. The total number of initially-loaded or filtered isolates, and the number of isolates shown in the graph (i.e. isolates with the metadata required for visualisation at the selected zoom levels) are summarized below the graph (as seen in **Supplementary Figure S3**).

The third interactive graph summarizes isolate counts over STs, CCs, or ODCs (example **Supplementary Figure S4**). It does not utilize any additional metadata, and only utilizes the MGT associated assignments. Isolate counts over STs, (or CCs, or ODCs) at each level are summarized in a table as a stacked bar chart. The interactivity enables a user to select any ST, CC or ODC value (by clicking on the corresponding bar), and view the distribution of isolates and breakdown of STs (or CCs or ODCs) at other MGT levels.

For each of these three visualisations, the user has to initially request the data by clicking on a button. Once requested, the entire data that may be required for subsequent plotting and switching-and-replotting is made available on the client's browser. The initial graph is generated and displayed at default zoom levels. The switching of a zoom level, filters and summarizes the data differently and refreshes the plot. We have observed that filtering is almost instantaneous, however, rendering of the stacked bar charts is slow, and this may be particularly evident when a user tries to view a large amount of data (such as all isolates in the database at the highest MGT level). To reduce render time, MGTdb collapses any STs (and CCs, and ODCs) assigned to a single isolate as a 'Singleton' class (depicted using white color in a stacked bar chart) when the number of isolates visualised are large (>1000 isolates) (example **Supplementary Figure S4**). This has improved performance, but may yet be slow when visualising the complete data on, for example *S.* Enteritidis, on computers with less working memory. In the future, we plan to explore further optimizations (e.g. by collapsing further STs into the singleton class, such as collapsing all STs containing ≤ 100 isolates when visualizing > 20000 isolates, i.e. 0.5% of total count).

To enable rapid and concise epidemiological investigations within countries, MGTdb provides summary reports (as for example, accessible here for *S.* Typhimurium https://mgtdb.unsw.edu.au/salmonella/summaryReport). The reports have been modelled to provide similar summaries as found in the Australian Department of Health's surveillance reports on notifiable enteric diseases[2]. A report on MGTdb aims to provide insights into recent trends by depicting isolate distributions of top STs via multiple charts. Once a country (or project) is chosen, the MGTdb web service retrieves data with the requested annotation within the specified range of years, from the organism database - then transforms the data for each chart and generates the plots, on the client's browser. Everytime a report is generated latest data is retrieved from the database, hence the report will always contain the most uptodate data with the requested annotations. Any generated report can be downloaded as an HTML document - thus, enabling the user to save a snapshot of the trends observed at a particular time. Finally, if a user is not logged in, the report is generated for publicly available data, and if a user is logged in, the report additionally includes isolates in the user's projects.

All charts in the report are static, i.e. cannot be interactively explored once generated. The report is systematically generated, and consists of the same charts, in the same order for any selected annotation. The initial charts in the report summarizes data to give an overview of the epidemiology. This is then followed by summaries at each MGT level. Four overview charts are plotted as follows (**Supplementary Figure S5**): the first depicts isolate counts per year, with the largest level MGT STs superimposed to enable a user to discern clonality in the population (**Supplementary Figure S5 A**), the second breaks down counts by sub-country or state - if the report is generated for a country, or breaks down isolates by country - if the report is generated for a project (**Supplementary Figure S5 B**). The third chart is a table that summarizes for all MGT levels, within the past three years (when user-specified range includes three years, otherwise the minimal range is used): the number of isolates, the number of STs and the number of STs newly observed in a particular year when compared to STs in the entire retrieved dataset (**Supplementary Figure S5 C**). The fourth chart depicts ST distributions for isolates at each MGT level (**Supplementary Figure S5 D**). Here, STs assigned to a single isolate are shown in white ('Singleton' class). These overview charts are followed by charts summarizing data at each individual MGT level. Six

---

[2] National Enteric Pathogens Surveillance System, Department of Health, Australia: https://www1.health.gov.au/internet/main/publishing.nsf/Content/cda-cdi3301r.htm#nepss (last visited on 2021, September 14).

charts are generated here, as follows: the first chart is a table that depicts the top 10 STs, along with isolate counts, percentages of isolates assigned each of the top 10 STs, and the clonal complex that the ST forms a part of, in the complete dataset. The second chart depicts the top 10 ST breakdown by year. The third chart presents the same information as in the second chart, but scales the ST distributions by the maximal ST distribution observed in that year. These two charts are seen in **Figure 3c**. The fourth chart breaks down the top 5 ST distribution by month (**Figure 3a**). The fifth chart consists of two sub-charts, which show for the most prevalent ST, the ST breakdown, at the previous and next MGT level for the last 3 years (**Figure 3b**). The last chart summarizes the top four STs distributed over the past four years (when user-specified range includes four years, otherwise the minimal range is used) as a bar chart.

In summary, MGTdb enables a number of different visualisation and data downloads, to enable effective epidemiological exploration at different levels of relatedness (such as closely related or distantly related) and for various types of users of the web-service (such as public health investigators or population geneticists).
